# Retrospective behavioral sampling (RBS): a method to effectively track the cognitive fluctuations driven by naturalistic stimulation

**DOI:** 10.1101/2022.05.30.493974

**Authors:** Talia Brandman, Rafael Malach, Erez Simony

## Abstract

Everyday experiences are dynamic, driving fluctuations across simultaneous cognitive processes. A key challenge in the study of naturalistic cognition is to disentangle the complexity of these dynamic processes, without altering the natural experience itself. Retrospective behavioral sampling (RBS) is a novel approach to track the cognitive fluctuations throughout the time-course of naturalistic stimulation, across a variety of cognitive dimensions. We tested the effectiveness and reliability of RBS in a web-based experiment, in which 53 participants viewed short movies and listened to a story, followed by retrospective reporting. Participants recalled their experience of 55 discrete events from the stimuli, rating their quality of memory, magnitude of surprise, intensity of negative and positive emotions, perceived importance, reflectivity state, and mental time travel. In addition, a subset of the original cohort re-rated their memory of events in a follow-up questionnaire. Results show highly replicable fluctuation patterns across distinct cognitive dimensions, thereby revealing a stimulus-driven experience that is substantially shared among individuals. Remarkably, memory ratings more than a week after stimulation resulted in an almost identical time-course of memorability as measured immediately following stimulation. In addition, idiosyncratic response patterns were preserved across different stimuli, indicating that RBS characterizes individual differences that are stimulus invariant. The current findings highlight the potential of RBS as a powerful tool for measuring dynamic processes of naturalistic cognition. We discuss the promising approach of matching RBS fluctuations with dynamic processes measured via other testing modalities, such as neuroimaging, to study the neural manifestations of naturalistic cognitive processing.

## 1 Introduction

Can we describe the sequence of cognitive processes engaged while watching a movie? Everyday experiences are dynamic, complex and multisensory in nature, supported by the interaction and regulation of simultaneously activated processing mechanisms (Hasson et al., 2004; Sonkusare et al., 2019). A fundamental challenge in the study of real-life cognition lies in tracking the cognitive processes that govern one’s continuously-changing experience of unfolding events. In recent years, a multitude of neuroimaging research groups has ventured beyond reductionist paradigms to mimic real-life experiences, such as watching a movie or listening to a story, while recording brain activation (e.g., Spiers and Maguire, 2007; Simony et al., 2016; Chen et al., 2017; Zadbood et al., 2017; Baldassano et al., 2018; Eickhoff et al., 2020; Brandman et al., 2021; Finn and Bandettini, 2021). Characterizing the fluctuations in cognitive states is key to understanding the fluctuations in neural and physiological states recorded during naturalistic stimulation.

There are three crucial problems in characterizing the cognitive dynamics driven by naturalistic stimuli as movies and stories. Unlike reductionist manipulations that isolate a specific target mechanism by onset (e.g., face perception: Farah et al., 1998; tone prediction: Bendixen et al., 2009), the multidimensional nature of naturalistic stimulation drives many simultaneous mechanisms of varying time-scales (Hasson et al., 2008). These can include a wide range of perceptual, semantic and emotional processes, social inference and memory encoding (e.g., McGurk and MacDonald, 1976; Loftus et al., 1978; Yeshurun et al., 2017). Furthermore, as information is continuously changing throughout unfolding events, we cannot predict how each of these simultaneous mechanisms will fluctuate throughout the movie. Thus, the first problem is disentangling the multitude of simultaneous processes, and the second problem is tracking their fluctuations across time. In addition, as we are interested in the natural, unaltered, human experience, our third problem is the act of measurement itself, which may affect in real-time the exact process we are trying to measure. This is widely known as the measurement problem, which, analogous to its origin in quantum physics, stipulates that “the mental content is invariably altered when the attention is concentrated on any special feature of it” (Bohr, 1958; Malach, 2007). How then, can we disentangle simultaneous cognitive processes activated during movie viewing, and measure their fluctuations along the time-course of the movie, without interfering with the natural ongoing experience?

To tackle these challenges posed by naturalistic stimulation, we developed a new approach termed retrospective behavioral sampling (RBS), which models the cognitive dynamics of human experience across a variety of cognitive dimensions. The central premise of this approach is based on previous work using autobiographical interview techniques (Berntsen, 2002; Levine et al., 2002), demonstrating that we can gain insight into one’s subjective past experience by reactivating their memory of it in retrospect. Such reactivation has been shown to evoke neural activity containing representations of the original experience (Gelbard-Sagiv et al., 2008; Chen et al., 2017; Zadbood et al., 2017; Norman et al., 2019). Following this logic, RBS is designed to reconstruct the cognitive state throughout the time-course of a dynamic stimulus, by deconstructing the experience into discrete events for reactivation. Following uninterrupted stimulation, the memory of each event is reactivated, and the (retrospective) subjective experience at the moment of the event is systematically reported across an array of cognitive dimensions.

We examined the effectiveness and reliability of RBS as a method for modeling cognitive dynamics driven by naturalistic stimuli such as movies and stories. The results presented below show that RBS patterns are informative and highly replicable across groups and testing sessions, and that individual differences are largely stable across stimuli. We assess the strengths and the limitations of RBS, and discuss its potential for matching cognitive processes with neural dynamics, individual clinical markers and other testing modalities.

## 2 Materials and Methods

### 2.1 Participants

Fifty-three participants were included in the study, in two groups: Group I included 26 participants (13 female, ages 33.08 ± 10.17) based in the USA. Group II included 27 participants (15 female, ages 34.48 ± 8.33) based in the UK. All participants were native English speakers with fluent writing abilities, and had intact hearing and eyesight (or corrected eyesight). Pre-screening further included a high approval record of at least 95% on previous web-surveys submitted on the survey platform, and technological compatibility criteria (see Experimental Apparatus). Two additional participants, not included above, had completed the experiment, but were excluded from analysis due to suspected cheating. This was determined by zero variance across measures on one or more events, reflecting repetition of the same rating response (usually “1”) throughout a long consecutive series of questions.

### 2.2 Stimuli

The stimuli for this study included four 4 short documentary movies and an auditory story. Two of the movies, ’Human Body’ (1:03 min; BBC) and ‘Pockets’ (3:08 min; Pilgrim Films), were adapted from the stimulus set of the Human Connectome Project (HCP; Cutting et al., 2012; Van Essen et al., 2013). The former was a narrated clip depicting people in extraordinary circumstances (e.g., a blind rock climber), and the latter was a street-interview clip in which passers-by presented and explained the content of their pockets. Two additional movies were narrated nature documentaries adapted from Planet Earth II (BBC), depicting Great Bowerbirds nesting and mating strategies (4:38 min), and Hyenas coexisting with humans in the city of Hara in Ethiopia (5:07 min). Finally, the auditory piece was a real-life story, ‘Pie Man’ (7:18 min; Jim O’Grady), recorded at a live storytelling performance (‘The Moth’ storytelling event, New York City), and adapted from the stimulus set used in earlier work (Simony et al., 2016).

### 2.3 Experimental Procedure

Experimental procedures were approved by the institutional review board (IRB; approval reference # 533-2) of the Weizmann Institute of Science.

Individuals who matched the pre-screening criteria on the web-survey platform were directed to an introductory page with a description of the task and technical compatibility requirements. Those who chose to continue and confirmed compatibility proceeded to the experiment. Participants then went through two additional screening steps to make sure that they could view the videos in their intended dimensions and that they could hear and see the stimuli well with their current setting (see Experimental Apparatus).

After passing all screening stages, participants were presented with the four videos in random order, and the auditory story presented last. Only after all stimuli were viewed participants were directed to the RBS questionnaire, in which they responded to each stimulus in turn, by order of initial stimulation. For each stimulus, participants first typed an open-ended free recall of the entire viewing experience, including all they could remember from the movie/story and their own reactions when watching. The free recall was followed by a series of rating questions corresponding to the same movie/story, in which each time, participants were reminded of a single event in the movie/story and rated how well they remembered it, understood it, how much it surprised them, the intensity of any positive and negative emotions it triggered, how important the event was to the main storyline of the movie/story, how much they were reflecting in thought versus immersed in stimulation at the beginning and at the end of the event, and how much the event prompted them to think about the past or the future. Altogether, participants completed 5 free-recall sessions, corresponding to the 5 stimuli, and 495 rating questions, corresponding to 9 measures X 55 sampled events.

Upon completing the RBS questionnaire, participants were asked to report their age and gender, and were asked whether they had been exposed to any of the 5 stimuli before participating in this experiment.

### 2.4 Experimental Apparatus

The experiment was conducted in an online web-survey written in HTML/CSS and JavaScript, and carried out on survey platforms Amazon Mechanical Turk (Group I) and Prolific.com (Group II). Technological compatibility screening included a computer operated with Windows, Mac, Linux or UNIX, Wifi connection sufficient for downloading ∼200 MB video content, working sound device, and a screen large enough to view the videos at fixed presentation size. Videos were presented at 200 mm × 112.5 mm, and participants were instructed to maintain a distance of about 30 cm from the screen. Screen size was validated in an interactive procedure, in which participants increased their browser window size until a button appeared allowing them to proceed with the experiment. Video presentation quality was validated with a practice video sample followed by catch questions to confirm participants both saw and heard the sample well. Prior test stimulation, participants were instructed to view every video and listen to the auditory story in full, without pausing, rewinding or skipping ahead. Pauses were recorded and viewing time was monitored to assure full and continuous viewing of each stimulus.

### 2.5 The RBS questionnaire

We developed the RBS questionnaire to track the temporal fluctuations in cognitive states along the stimuli timeline. To achieve this, we densely sampled events along the stimulus timeline at intervals of around 15 to 30 seconds. The main consideration for selection of events within these interval timeframes was that they could be coherently described, i.e., that something distinct was happening that could be used as a reminder to recall the event. Throughout the 4 movie clips, 33 events were sampled, and 22 more events were sampled throughout the auditory story, resulting in a total of 55 events.

The questionnaire was blocked by stimulus, such that after viewing all stimuli consecutively, participants completed the entire RBS questionnaire by order of stimulus viewing. Each stimulus block began with a typed free recall of the entire movie or story, including the subjective experience of the participant while viewing. After free recall, participants received instructions for event rating, in which they were asked to refer specifically to each event, meaning no more than a few seconds before or after the described moment in the movie/story. General instructions for the event rating phase were given as follows: “In the next pages, you will be prompted on particular events in the movie/story. Each time, you will receive a description of an event. For example: *narrator:* “*Lions can see well in the moonlight. This one is on the lookout for prey"*. (1) You will be asked to recall the precise moment in the movie/story - no more than a few seconds before and after this event (not the entire scene) - and rate the quality of your memory of it. (2) You will be asked to answer a series of questions about your experience in that moment of the movie/story. Read the instructions of each question carefully, and use the full range of the scale between 1-7 to provide a response that most accurately describes your experience, as you remember it, at the time of watching/hearing the particular event in question. The answers should describe your personal experience while watching/listening, as accurately as you can remember".

Following these general instructions for event rating, participants began the event-rating phase of the block. Each time participants would be probed on a single event from the movie/story and rate it on all tested measures in a sequence of multiple-choice questions. All sampled events for the movie/story were rated. Events were probed in random order.

Each event began with a textual reminder. For example: *Mustached man eating little sweets:* “*Yes it‘s salty, and*… *it’s licorice”* (all event reminders are listed in Supplementary Table 1). This screen appeared for 30 seconds, in which participants were instructed to silently recall the event in their mind, including everything they could remember about what happened and their own subjective viewing experience. The exact instructions were: “Please take a moment to recall this exact moment of the movie, what happened, what you saw, what you heard, what were your own thoughts, emotions and/or physical sensations at that moment, etc.". The objective of the silent recall was to bring the earlier experience of viewing the event into mind. When the time was up, the screen automatically changed to the first rating question.

Each rating question appeared in single page, with the event reminder kept at the top of the page during the entire rating sequence. No time limit was given for response. Upon response (key press between 1 and 7), participants were automatically forwarded to the next question.

#### 2.5.1. Rating questions

##### Memory

before rating any specific type of experience, participants were asked to rate their quality and detail of memory for the recalled event. The exact phrasing was: “How would you rate your quality of memory for this event as compared to other events? Consider the full range of the scale and provide an accurate response between 1 – no memory at all – to 7 – best and most detailed remembered event in this video".

##### Comprehension sanity check

“How would you rate your comprehension of this event? Consider the full range of the scale and provide an accurate response between 1 – did not understand what was happening – to 7 – understood perfectly what was happening".

##### Surprise

“How surprising was the event? Consider the full range of the scale and provide an accurate response between 1 – did not surprise me at all – to 7 – no other event in this video surprised me this much".

##### Negative Emotion

“How much negative emotion did this event cause you? Consider the full range of the scale and provide an accurate response between 1 – no detectable negative emotion – to 7 – more intensely negative than any other in this video".

##### Positive Emotion

“How much positive emotion did this event cause you? Consider the full range of the scale and provide an accurate response between 1 – no detectable positive emotion – to 7 – more intensely positive than any other in this video”

##### Importance

“How important was this event for understanding the main storyline of this video? Consider the full range of the scale and provide an accurate response between 1 – insignificant – to 7 more important than any other event in this video".

##### Initial Reflectivity

“When this event came up, at what state of mind did it “catch” you? How much were you reflecting on things either related or unrelated to the movie immediately before this event happened? (such as recounting previous events or personal memories, predicting what would happen next, analyzing the plot or characters, or any other type of thought). Consider the full range of the scale and provide an accurate response between 1 – completely immersed in video, no other thought – to 7 most reflective moment in this video".

##### Triggered Reflectivity

“How did this event affect your state of mind? How much did this event make you reflect on things either related or unrelated to the movie immediately after it happened? (such as recounting previous events or personal memories, predicting what would happen next, analyzing the plot or characters, or any other type of thought). Consider the full range of the scale and provide an accurate response between 1 – completely immersed in video, no other thought – to 7 – most reflective moment in this video".

##### Mental Time Travel

“What did this event make you think of - did it send your thoughts into the past or into the future? In other words, how much did it remind you of past events or make you predict what would happen next? Consider the full range of the scale and provide an accurate response between 1 – vivid recollection of past events – to 7 – vivid prediction of future events".

### 2.6 Follow-up testing

A subset of 23 participants from Group I re-rated their memory of all events between a week to two weeks following original testing. During follow-up testing, participants did not view the stimuli again, and were directly taken to a stripped-down version of the RBS questionnaire. In each block, they completed the typed free recall of the entire stimulus, and the silent recall and memory rating for every event, as in the original questionnaire, but were not asked any further questions.

### 2.7 Data processing

Trial exclusion: In the case that a participant had reported previous exposure to a stimulus, i.e., that the experiment had not been their first viewing of a certain movie/story, only the events corresponding to that stimulus were excluded from that participant’s data. Previous exposure was reported for the Bowerbird video in 3 participants and for the Hyena video in 6 participants. In addition, in the case that a participant presented poor understanding of a certain stimulus, all events corresponding to that stimulus were excluded from their data. Particularly, one participant rated their comprehension between 1-3 (out of 7) for 16 out of the 22 events in the auditory story, whereas the same participant’s comprehension ratings for the movie events were high. We therefore excluded the story events from this participant’s results.

Sanity checks: Following trial exclusion as described above, comprehension ratings were examined as a sanity check, to ensure overall coherency of sampled events and their textual reminders. Means across subjects of event comprehension suggest that all events and probes were sufficiently coherent, with comprehension of 5.74 (out of 7) on average across all events, and the lowest mean comprehension of 4.83 on Event #9.

Scoring: To prepare rating data for statistical testing, z-scoring was performed along the entire dataset of each participant separately, by subtracting the mean and dividing by standard deviation.

Individual Typicality: The typicality index for each participant was defined by the following equation, as the Euclidean distance (*d*) between the feature vector of each participant (*q*) and that of the group (*p*), given by the square root of the sum of squared deltas for each element *i* of *q* and *p* between 1 and *n*. Euclidean distance was scaled by the square root of vector length (*n*).

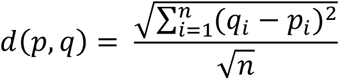

## 3 Results

RBS began with uninterrupted stimulation using movies or stories, followed by a behavioral questionnaire collecting a systematic report of the cognitive experience corresponding to each event in the movie. The details of the approach are presented in Figure 1. To reactivate one’s experience of a specific event, it was described by a textual reminder, with time taken to silently recall all details of the event and the subjective experience it had evoked. Thereafter, the experience was rated on a series of cognitive scales measuring the quality of memory for the event, how surprising it was at the time, the intensity of positive and negative emotions triggered by the event, and perceived importance of the event to the main storyline. In addition, introspective rating questions were aimed at modeling one’s state-of-mind around the time of the event. These included the extent to which one felt immersed in the movie or reflective in thought, at the time leading into the event and later as triggered by the event. Finally, mental time-travel referred to the extent to which such reflections concerned the past or the future.

**Figure 1.**
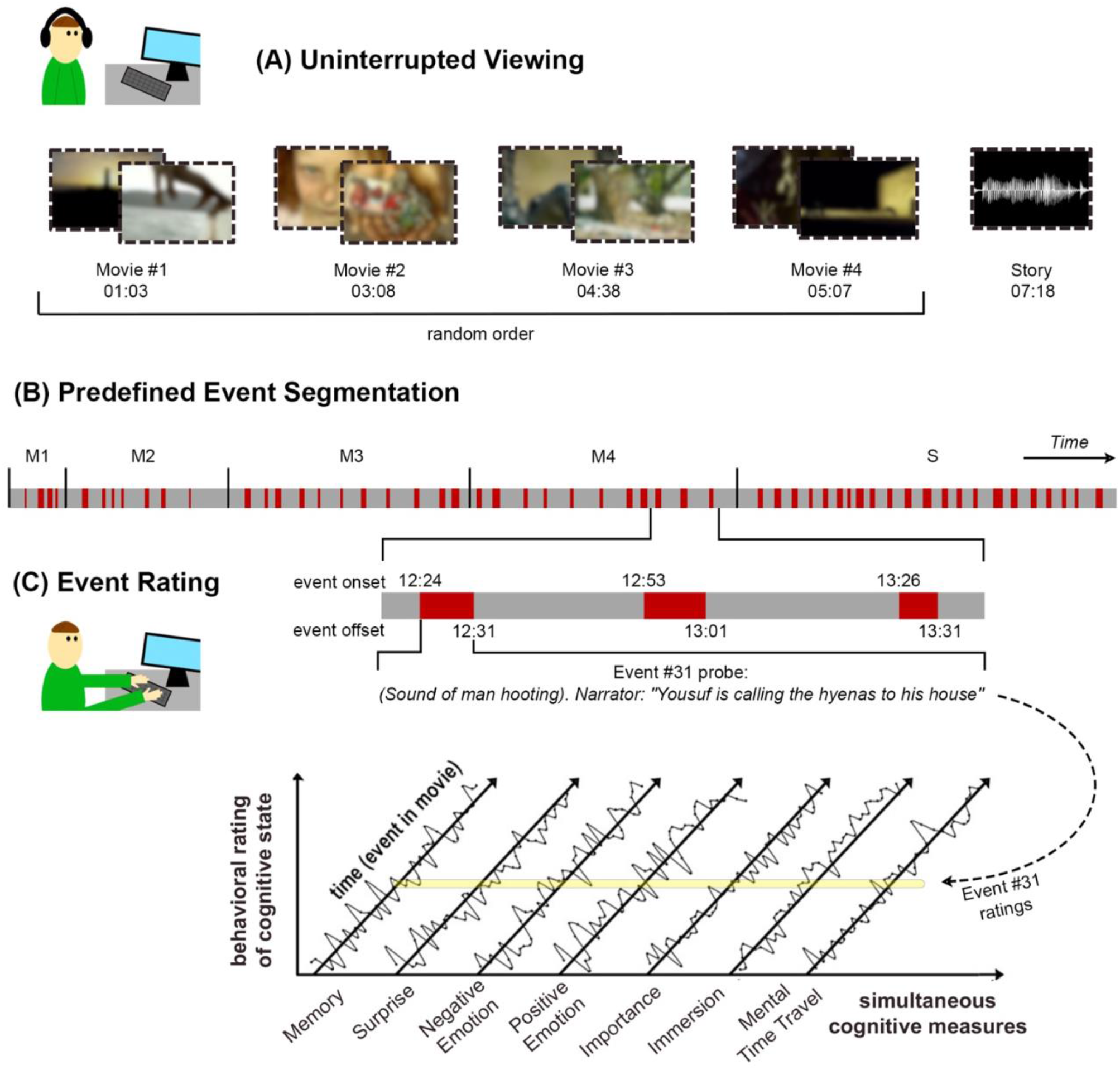
Experimental approach. (A) In the first part of the task, participants viewed 4 short movies (in random order) and listened to a story, sequentially with no interruptions. The duration of each stimulus is detailed as min:sec. The stimuli were streamed online to participants’ personal devices. (B) Prior the experiment, textual descriptions were generated to match a total of 55 stimulus events, with 4 events in Movie #1 (M1), 7 in Movie #2 (M2), 11 for each of Movies #3 (M3) and #4 (M4), and 22 events for the story (S). Event onset and offset (as min:sec) match the beginning and end of the happenings described in the textual event reminder. (C) In the second part of the task, participants filled out the retrospective behavioral sampling (RBS) questionnaire, in which they viewed the textual reminder for each event (in random order within each stimulus block) and rated their experience of it on an array of cognitive dimensions.

### 3.1 How effective is RBS for dynamic cognitive tracking?

We set out to assess the efficacy of RBS as a reliable and informative method for cognitive tracking of dynamic natural experience. To this end, we tested the replicability of the sampled cognitive dynamics across independent groups and testing sessions, as well as their stability by group size, and informativeness across time.

First, we examined whether RBS is generally informative of differences between events and between measures. Indeed, we found significant differences between sampled events, which further varied as a function of cognitive measure. This was tested in repeated-measures ANOVA of ratings, with event (1 through 55) and cognitive measure (memory, surprise, negative emotion, positive emotion, importance, initial reflectivity, triggered reflectivity, mental time travel) as within-subject factors, which revealed main effects of event (F(54,2430) = 19.23, P< 0.001), and measure (F(7, 315) = 105.99, P< 0.001), as well as an interaction between them (F(378,1.7*e*^4^) = 11.68, P< 0.001).

Next, we asked whether these cognitive dynamics, given by the time-courses of ratings across events, can be replicated across the two independent groups of participants (N_1_=26, N_2_=27). Correlation analysis revealed robust replicability of cognitive dynamics across groups in all tested measures (FDR corrected P< 0.001). Specifically, mean ratings were correlated across the 55 events between the two groups at R = 0.9 for Importance, Positive Emotion, Negative Emotion, and Surprise, R = 0.8 for Memory, R = 0.7 for Triggered Reflectivity and Mental Time Travel, and R = 0.5 for Initial Reflectivity. Detailed correlation values are presented in Figure 2a.

**Figure 2.**
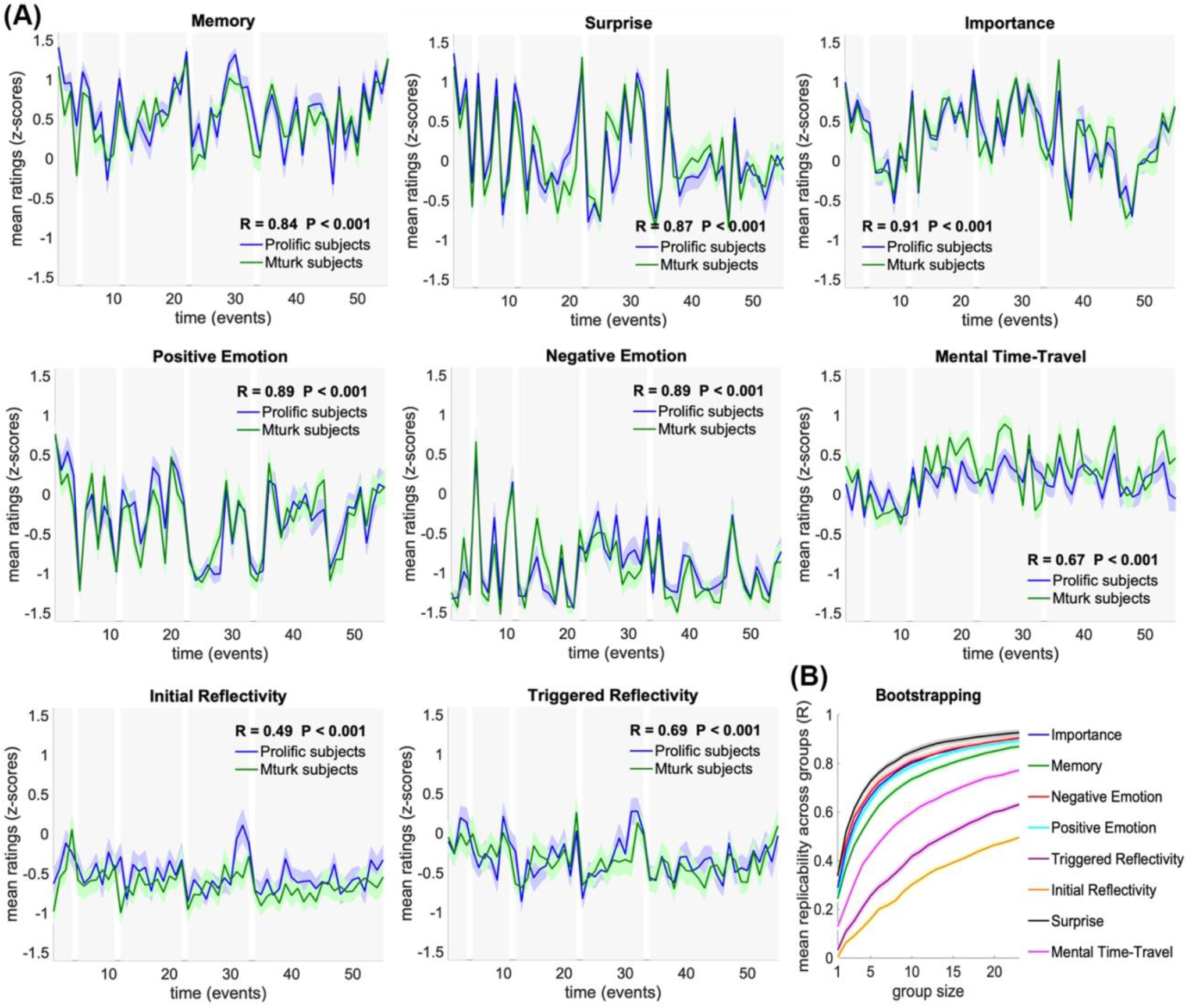
Replicability and stability of cognitive dynamics. (A) Comparison of mean ratings across subjects between Group I - Mturk subjects (green; N=26), and Group II - Prolific subjects (blue; N=27). Dynamics were significantly correlated between groups in all tested cognitive measures (FDR corrected P< 0.001). Ratings are presented as mean and SEM across subjects. (B) Stability of group dynamics as a function of group size in a 100-iterations bootstrapping procedure. Each iteration randomly assigned 23 participants to each of 2 independent groups and tested the correlation across group means for each group size between 1 and 23. Bootstrapping results are presented as mean and SEM across sampling iterations.

To further examine the stability of these results as a function of group size, we calculated the correlations of randomly assigned groups sampled 100 times for each group size, ranging from 1 to 23 participants per group (of the 46 participants with full trial data). This bootstrap analysis revealed high stability, with mean correlations reaching R>0.4 with as few as 3 participants for Surprise, Memory, Importance, Negative Emotion and Positive Emotion, and with 4 participants for Mental Time Travel, relative to moderate stability of Initial and Triggered reflectivity, which reached R>0.4 with 17 and 10 participants, respectively. Detailed correlation values are presented in Figure 2b.

Finally, to assess test-retest reliability of RBS, we examined the replicability of memory ratings in a subset of participants (N=23) who had rated their memory of all events a second time, between a week to two weeks after stimulation. We found strikingly high replicability across the two rating sessions, with mean ratings of Memory correlated across the 55 events between the two testing sessions at R = 0.9 (P< 0.001) (Figure 3). We note that, while this confirms test-retest reliability, it does not rule out potential effects of initial reporting on memory ratings in the follow-up session.

**Figure 3.**
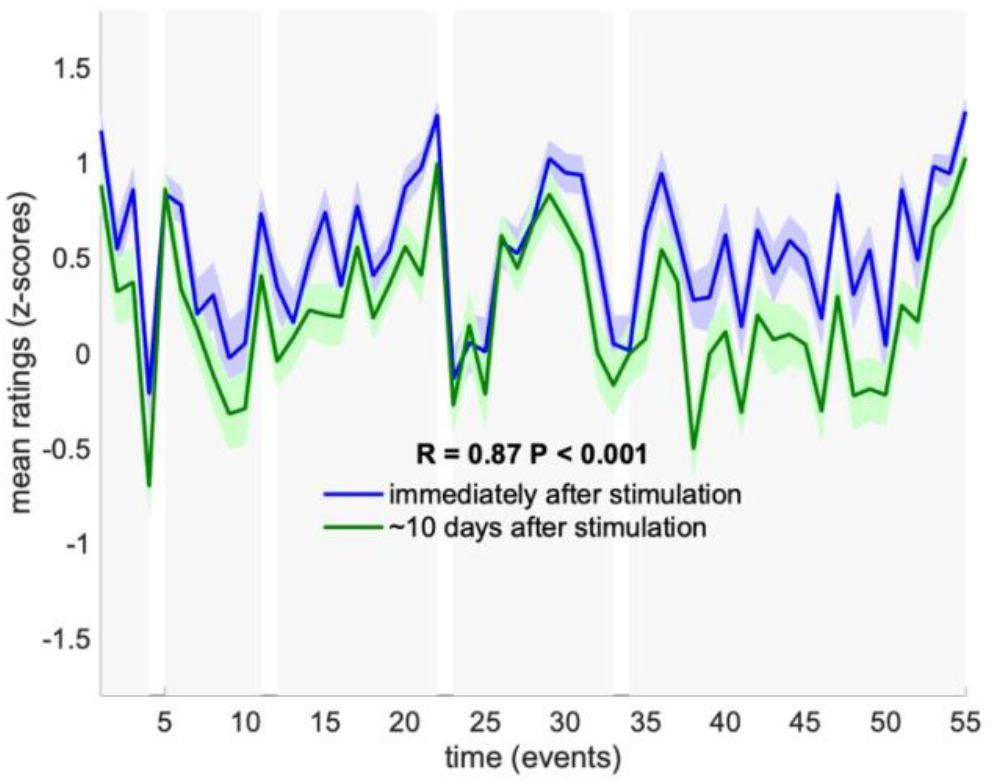
Test-retest reliability of memory ratings. Mean memory ratings across subjects who participated in the follow-up test (N=23), were compared between the first rating session immediately after stimulation (blue) and second rating session ∼10 days later (green). Dynamics were significantly correlated across rating sessions (P< 0.001). Ratings are presented as mean and SEM

### 3.2 How are cognitive measures related to one another?

To examine the relationship among the different cognitive measures, we constructed their pair-wise correlation matrix for each participant, by correlating across the 55 events between each pair of cognitive measures. The group means of within-subject correlations are presented in Figure 4a. In addition, we calculated the correlations between each pair of measures across the two independent groups of participants, such that each measure in one group was correlated with the second measure in the second group. The symmetrized matrix of correlations between groups is presented in Figure 4b. In both correlation matrices, the four largest collinearities among cognitive measures were observed between Initial Reflectivity and Triggered Reflectivity, Positive and Negative Emotion (inverse), Memory and Importance, and Memory and Surprise (FDR corrected P< 0.001).

**Figure 4.**
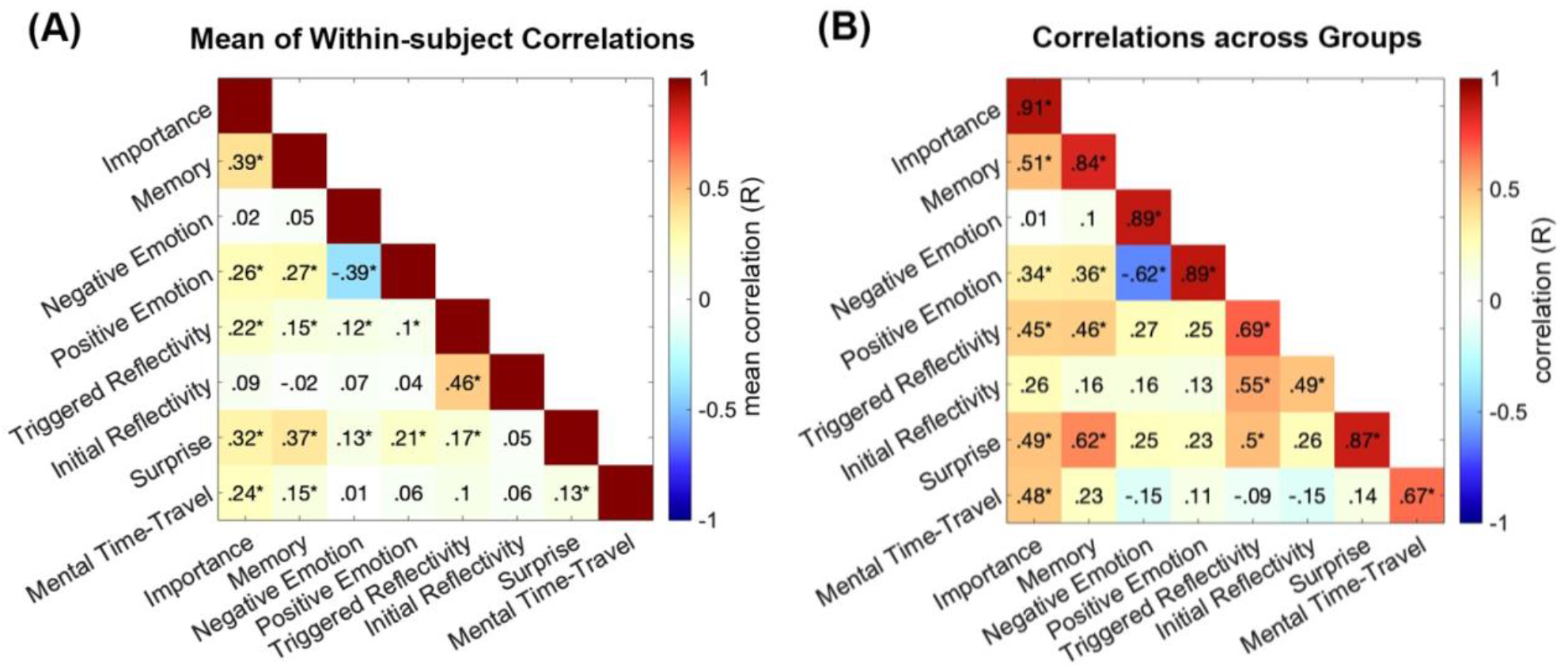
Collinearities among cognitive measures. Correlations were calculated between each pair of cognitive measures across all 55 events. (A) Presents the mean across subjects of correlations calculated for each individual participant. Significance for each pair of measures was assessed in a t-test of the Fisher-transformed correlation coefficients, across subjects, against zero. (B) Presents the symmetrized matrix of correlations between the two groups. Significance for each pair was tested on its mean correlation coefficient. * FDR corrected P< 0.05.

These collinearities raise the question of redundancy, i.e., whether any one measure could be fully explained by its shared variance with other measures. To test this, we computed the partial correlations across measures and groups. This was done by correlating each single measure in Group I with the same measure in Group II, while controlling for correlations between Group I and all other measures in Group II. Partial correlations were calculated in both directions, i.e., between Group I and II while controlling for all other measures in Group II and vice versa, then averaged across the two results. Mean partial correlations: Importance R = 0.83 (FDR corrected P< 0.001), Surprise R = 0.79 (cor. P< 0.001), Memory R = 0.73 (cor. P< 0.001), Positive Emotion R = 0.66 (cor. P< 0.001), Negative Emotion R = 0.64 (cor. P< 0.001), Mental Time Travel R = 0.47 (cor. P = 0.002), Initial Reflectivity R = 0.19 (cor. P = 0.596) and Triggered Reflectivity R = 0.17 (cor. P = 0.674). Thus, only collinearities among measures of Reflectivity (both initial and triggered) explained the full variance driving replicability of these two measures across groups.

### 3.3 How effective is RBS for studying individual differences?

To test the potential use of RBS for the study of individual differences, we calculated participants’ (a)typicality index, defined as the scaled Euclidean distance between his or her rating pattern and that of the group mean (Figure 5a) (Ramot et al., 2020). Thus, the typicality index provides an assessment of how different an individual is in his or her unique pattern of response relative to the group. For example, a rating time-course similar to the group mean would be indexed as typical – given by a *low* typicality index, whereas a rating pattern more different from the group mean would be indexed as less typical – given by a *high* typicality index (Figure 5b). We note that the results reported below did not change when, instead of using the full group mean, typicality scores were calculated against the group mean excluding the target subject.

**Figure 5.**
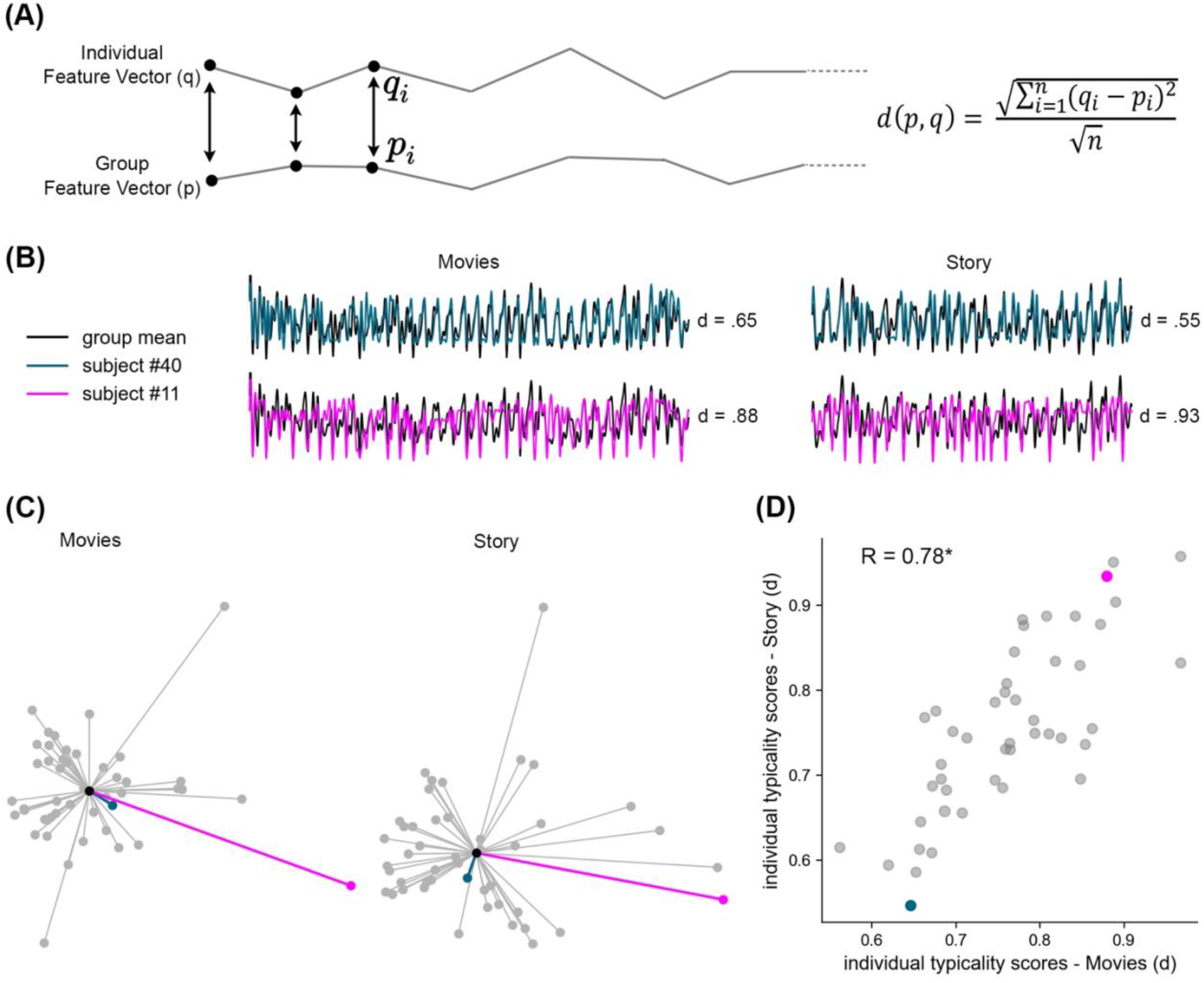
Individual typicality of cognitive dynamics. (A) Individual typicality was measured by the Euclidean distance (*d*) between the feature vector of each participant (*q*) and that of the group (*p*), given by the square root of the sum of squared deltas for each element *i* of *q* and *p* between 1 and *n*. Euclidean distance was scaled by the square root of vector length. Individual typicality was separately measured across the 33 movie events and across the 22 story events, over 8 cognitive measures (movies n = 264, story n = 176). Panels B through D illustrate the stability of individual typicality across movies and story, highlighting two example participants, #40 (cyan) and #11 (pink), relative to the group mean (black). (B) Rating dynamics of the two example participants overlaid on the z-scored group means, and their typicality indices. (C) Illustration of the distances between every participant and the group mean, under Isomap 2D dimensionality reduction. Each node represents a single participant and each edge represents the participant’s Euclidean distance from the group mean (center node). (D) Individual typicality scores, calculated in full dimensionality, were significantly correlated across subjects between movies (X axis) and story (Y axis). Each data point represents a single participant. * P< 0.001.

To test whether individual typicality captured by RBS dynamics is a reliable marker of individual differences, we asked whether it is stimulus-invariant, i.e., whether it would be stable across different types of narrative stimuli - even when they differ in content and modality. To test this, we calculated the typicality index for movies (33 events) and, separately, the typicality index for the auditory story (22 events), for each participant with complete trial data (N=46). The distribution of typicality indices is illustrated in Figure 5c in reduced dimensionality. We then measured the correlation across participants’ typicality indices between the two types of stimuli, revealing a robust correlation of R = 0.78 (P< 0.001), which thereby explains 61% of the variance across individuals (R^2^ = 0.61). As visible in Figure 5d, all participants showed stable typicality across stimuli.

While the above typicality analysis harnessed the full variability of the temporal response patterns across individuals, we additionally calculated the specific typicality indices corresponding to each cognitive measure separately (see Supplementary Figure 1). Correlations across stimuli varied by measure: Mental Time Travel R=0.77 (FDR corrected P< 0.001), Memory R = 0.75 (cor. P< 0.001), Negative Emotion R = 0.72 (cor. P< 0.001), Triggered Reflectivity R = 0.62 (cor. P< 0.001), Initial Reflectivity R = 0.62 (cor. P< 0.001), Positive Emotion R = 0.49 (corrected P< 0.001), Importance R = 0.18 (cor. P = 0.255), Surprise R = 0.10 (corrected P = 0.498).

## 4 Discussion

RBS was developed as a novel approach to reconstruct the fluctuations in cognitive states driven by naturalistic stimuli such as movies and stories. With RBS, we aimed to disentangle simultaneous cognitive processes, and model their fluctuations along the time-course of stimulation, without interfering with the direct natural experience. Results clearly show that RBS is both effective and reliable in characterizing these cognitive fluctuations. RBS captured changes in the cognitive state across events, which differed as a function of cognitive measure. These cognitive dynamics were highly replicable across independent groups of participants, indicating a substantial and reliable stimulus-driven experience. RBS was highly effective in tracking these shared dynamics, exhibiting striking replicability even with small sample sizes, as well as across separate testing sessions. Furthermore, RBS was informative of individual differences in subjective experience, showing consistency of individual typicality across different narratives and stimulus modalities.

### 4.1 Disentangling simultaneous cognitive processes

Cognitive dynamics, measured by RBS rating time-courses, varied across the 8 tested cognitive measures, consistently beyond individual variability. Furthermore, the specific dynamics corresponding to each cognitive process, separately, exhibited robust reliability. Together, these results suggest that RBS can effectively dissociate the temporal dynamics of several cognitive processes within a single questionnaire. Note, however, that the ability to dissociate these processes does not rule out the existence of collinearities among them, as well as non-linear interactions. In the current data, we see the largest collinearities among memory, surprise and importance – consistent with concepts of predictive processing, i.e., that deviance from expectation strengthens encoding (Rescorla and Wagner, 1972; Schultz, 1998), as well as between positive and negative emotion, and between initial and triggered reflectivity. Nevertheless, after controlling for all collinearities, the resultant unique component of each measure that is replicable across groups, remained high for all measures other than reflectivity. This suggests that the two reflectivity measures could be collapsed into a single question in future, with no expected loss of information.

### 4.2 Measuring stimulus-driven cognitive dynamics

As we have seen, rating time-courses exhibited robust replicability across independent groups of participants, suggesting that stimulus-driven experience encompasses typical patterns of cognitive dynamics that are shared across individuals. This can be viewed as analogous to the shared component of response found in stimulus-driven neural fluctuations, when correlating neuroimaging responses across individuals using dynamic inter-subject correlation techniques (Hasson et al., 2004; Simony et al., 2016). Such shared neural dynamics have been previously shown to reflect high-level cognitive representations that are shared across individuals (e.g., narrative interpretation: Nguyen et al., 2019; social inference: Ramot et al., 2020). In the current study, the shared element of cognitive experience was found to be effectively and reliably measures by RBS along the stimuli timeline, with minimal sample size. This makes RBS an ideal tool to generate predictor variables that can be applied to the analysis of fluctuations in a variety of testing modalities (Simony and Chang, 2020). In essence, using RBS, we can associate cognitive processes with any other dynamic process corresponding to the same stimulus, e.g., as measured with neuroimaging, electrophysiology, dynamic physiological indices or eye-tracking.

Showcasing its potential, our previous work using an earlier (provisional) version of RBS, uncovered correlations between default mode network (DMN) coactivations and experienced surprise, pointing to an important role of the DMN in predictive processing of movie narratives (Brandman et al., 2021). The discovery was made possible by comparing the RBS surprise ratings collected in a web-based experiment, with neural dynamics from earlier functional magnetic resonance imaging (fMRI) studies (Chen et al., 2017; Zadbood et al., 2017), thereby matching cognitive and neural fluctuations across independent groups of participants. This highlights an important advantage of RBS, namely, that it does not require that the same participants undergo both RBS and neuroimaging. Thus, RBS can be used to extract cognitive regressors for an entirely independent dataset, allowing for its immediate implementation to investigate the cognitive processes reflected in brain signals recorded in earlier neuroimaging and electrophysiological studies. The only prerequisite is that RBS be collected for the same stimuli as in the dataset to be matched. In addition, using a similar approach, we could compare the dynamics of human experience, as measured with RBS, with computer inference, as given by artificial neural-network output to the same stimuli.

### 4.3 Individual differences in cognitive dynamics

RBS rating patterns were indicative of the extent to which an individual’s reported experience was typical relative to the common experience of the group. Individual typicality was found to be stable across movies and story stimuli, thereby tapping into stimulus-invariant individual response tendencies. These results highlight the functional power of coding through similarity distances, which have been previously shown to explain perceptual differences as well as brain specializations (Davidesco et al., 2014; O’Connor et al., 2017; Finn et al., 2020; Ramot et al., 2020; Levakov et al., 2021; Malach, 2021).

When examining each cognitive measure on its own, we found stimulus-invariant typicality for all but surprise and importance, yet no clear relationship or trade-off was observed between stability of individual typicality and the robustness of group dynamics (i.e., replicability). At this point, we refrain from inferring about long-term cognitive traits from these data, as further research will be needed to assess whether RBS typicality is informative of stable trait information beyond a single session (i.e., by splitting stimulus sets by testing days).

The current findings call for two potential approaches to examine individual differences utilizing RBS individual typicality. The first approach is applicable when neural (or physiological) responses are recorded during sensory stimulation, e.g., movie viewing, prior to applying RBS to the same stimuli. In this case, RBS can be used to match individual cognitive typicality to its corresponding neural typicality for the same movie, such as the distance between the participant and the group in the activation of a region of interest (inter-subject correlation: Hasson et al., 2004), or its coactivation with another region (Simony et al., 2016; Simony and Chang, 2020). This type of comparison does not rely on stimulus-invariant information, and thus can potentially relate cognitive processes that lack that invariance to simultaneous neural processes, in a manner that reflects subjective fluctuations in the neurocognitive state. Thereby, RBS typicality can be used to explore the neural mechanisms related to subjective processing of specific cognitive dimensions, which may be canceled-out in the common group dynamics. The second potential application of RBS typicality extends to stimulus-invariant information. In this case, RBS typicality, being a special form of representational distance measure, could be matched with any marker of individual differences, in any modality, regardless of stimulation (e.g., Grossman et al., 2019). For example, future research could examine the diagnostic value of behavioral naturalistic-stimulation settings, by comparing RBS typicality with performance scores on clinical questionnaires or tasks. A similar approach using representational distances has been successfully applied in previous clinical research (Hahamy et al., 2015).

### 4.4 The measurement trade-off: real-time versus retrospective reporting

RBS was designed to reconstruct the cognitive processes evoked by naturalistic stimulation, without interfering with the natural experience as it is happening. Thereby, we avoid the measurement problem, by which the introspective attention and reflection upon one’s own experience, required for explicit reporting in real time, risks altering the experience itself (Bohr, 1958; Goldberg et al., 2006; Malach, 2007). Instead of interrupting the natural experience, retrospective reporting relies on one’s memory of it. Particularly, RBS was inspired by previous work showing that one’s subjective past experience can be reactivated through memory, in great detail that goes beyond episodic recall of the facts to include emotional and introspective perceptions (Berntsen, 2002; Levine et al., 2002). Accordingly, in RBS, the experience of each event is first reactivated in a silent-recall stage, followed by the rating of subjective experience. In addition, a practical advantage of RBS over real-time reporting is the wider scope of information that can be recovered. Unlike real-time reporting, which is limited in the number of simultaneous processes to which one can attend and report on at a given moment, RBS probes a variety of distinct processes that are each recalled from memory in turn.

On the other hand, the reliance on memory for retrospective reporting comes at a price, namely, that we are measuring the memory of an experience, not the experience itself. Hence, in RBS, positive emotion is the memory of feeling good, and surprise is the memory of being surprised, etc. The gap between experience and the memory of it has been the main rational for advocates of real-time rather than retrospective paradigms (Kahneman and Riis, 2005; Schwarz, 2007). By this logic, in the lack of an associated real-time measure, we cannot disentangle RBS real-time validity from memory distortion. Yet RBS was developed with the idea in mind of matching retrospective reports with real-time neuroimaging dynamics. Critically, when matching RBS with any real-time recorded measure, such as neuroimaging, their shared variance across events will represent a real-time component of experience. In this approach, the resultant real-time component will not be contaminated by interruptions to stimulation. By contrast, real-time cognitive reporting by neuroimaging participants will require interference with stimulation during neuroimaging recording, thereby contaminating both measurement modalities in a way that is irrecoverable. We therefore argue that, considering all aspects of the measurement trade-off, RBS is beneficial for achieving high validity of neurocognitive processes, while maximizing the ecological nature of real-time experience.

### 4.5 Conclusions

RBS is shown to be an informative and reliable method to track natural cognitive dynamics, successfully tackling the central challenges of naturalistic stimulation; RBS effectively disentangles a wide range of simultaneous cognitive processes, depicts them along the time-course of stimulation, and does this without interfering with the real-time experience. The robust replicability of RBS time-courses makes it a powerful tool to characterize the shared stimulus-driven cognitive experience. These shared dynamics can be applied to ascribe specific cognitive processes to neural and physiological time-courses driven by the same stimuli. RBS was further shown to carry stimulus-invariant information on individual differences, which could be tested against neural and clinical individual markers to study idiosyncratic neurocognitive states and traits. In sum, RBS is a promising new tool to advance the study of naturalistic cognition and its neural and physiological manifestations.

## 6 Author Contributions

Conceptualization and writing by T.B., R.M., and E.S., funding acquisition by E.S., experimental design, data collection, data analysis and visualization by T.B.

## 8 Acknowledgments

We thank Sharon Pinto and Ronen Pinto for web development supporting the survey distribution and data aggregation and export.

