## Supplementary for "Retrospective behavioral sampling (RBS): a method to effectively track the cognitive fluctuations driven by naturalistic stimulation"

**Supplementary material for preprint:** *Retrospective behavioral sampling (RBS): A method to effectively track the cognitive fluctuations driven by naturalistic stimulation*

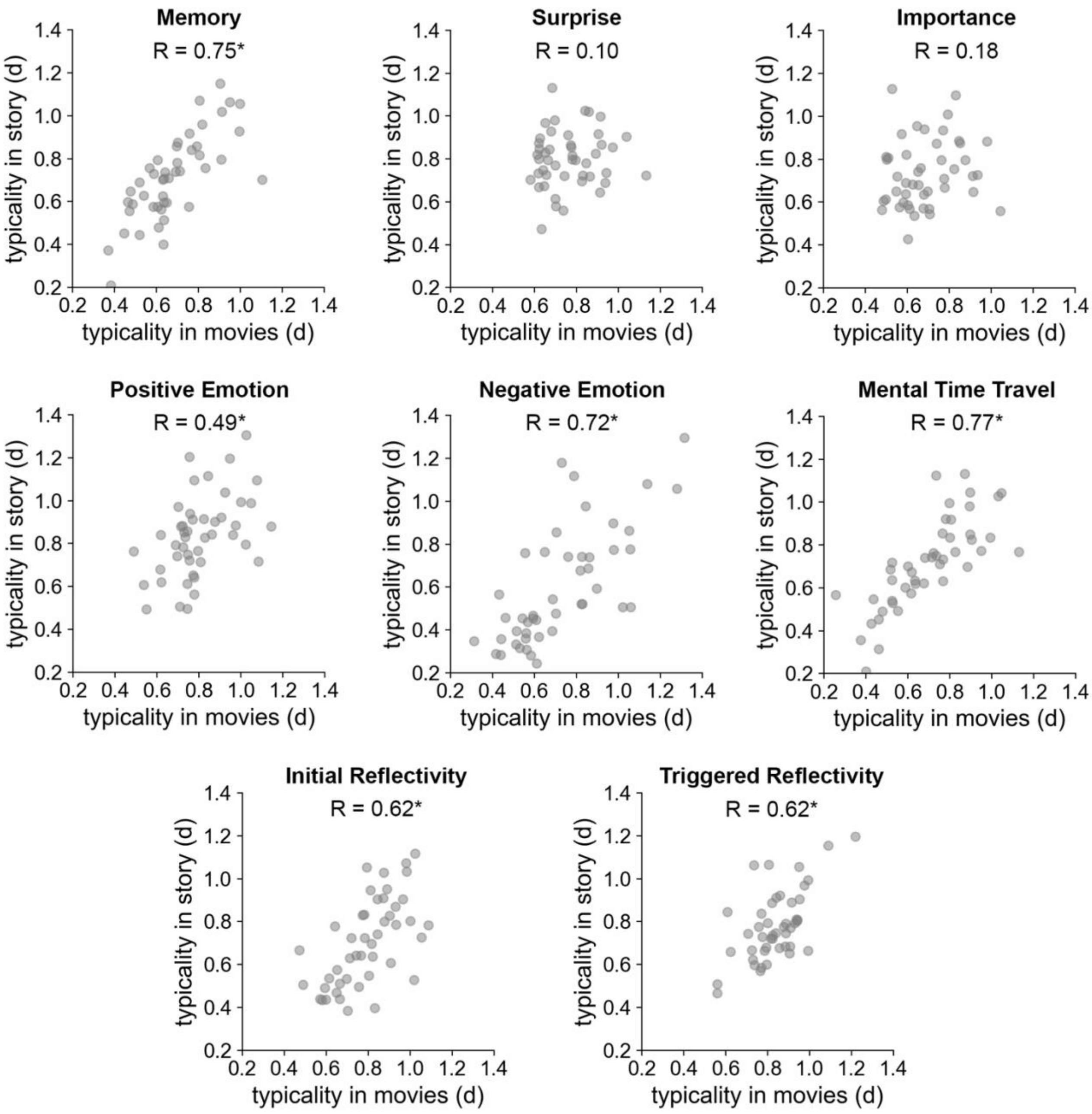


**Supplementary Figure 1.** Individual typicality within each cognitive measure. In all measures except for Surprise and Importance, individual typicality scores were significantly correlated between movies (X axis) and story (Y axis). Each data point represents a single participant. * P < 0.001.

| # | Movie | Textual Reminder |
|---|---|---|
| *1* | Movie 1: ‘Human body’ | 0:19 - Man climbs cliff. Narrator: "This man is blind" |
| *2* | | 0:34 - Narrator: "These fishermen can see underwater, better than almost anyone else on Earth" |
| *3* | | 0:45 - Narrator: "And this delightful young girl is alive an well, despite having just half a brain" |
| *4* | | 0:54 - Narrator: "Their stories are part of your story" |
| *5* | Movie 2: ‘Pockets’ | 0:22 - Young woman: "In my pocket I've got a crack pipe, cigarettes and money, and that's what I use every day" |
| *6* | | 0:45 - Moustached man eating little sweets: "Yes it's salty, and... it's licorice" |
| *7* | | 0:56 - Young boy with key: "The key opens nothing" |
| *8* | | 1:07 - Old woman with ring: "And in the end I had to offer him sex to get it off of him" |
| *9* | | 1:34 - Old man feeds dog from hand |
| *10* | | 1:53 - Young man with memorial card: "I was only young when she died, but she was always there for me" |
| *11* | | 2:25 - Young man with medical printout: "Last week I made love with like five girls" |
| *12* | Movie 3: ‘Bowerbirds’ | 00:21 - Narrator: "This great bowerbird has spent over a decade building this collection of mostly man-made objects" |
| *13* | | 00:44 - Bird moves a piece of styrofoam. Narrator: "Perhaps that would look a little better over there" |
| *14* | | 00:56 - Narrator: "Instead of going into town to collect new objects, he's decided to raid his neighbour's bower" |
| *15* | | 01:24 - Narrator: "The owner is back". Bowerbirds screech and hiss at each other. |
| *16* | | 01:45 - Narrator: "He'll have to wait for the owner to leave" |
| *17* | | 02:11 - Bowerbird first steals scarlet heart. Narrator: "Got it!" |
| *18* | | 02:35 - Guest Bowerbird arrives. Narrator: "The seduction can now begin" |
| *19* | | 03:04 - Narrator: "But his guest doesn't seem to be paying much attention" |
| *20* | | 03:36 - Narrator: "But he still has one trick up his sleeve. The scarlet heart" |
| *21* | | 04:05 - Narrator: "As a final thrill, he expands the pink crest on the back of his head" |
| *22* | | 04:19 - Narrator: "This is not a female but a young male who hasn't yet developed that head crest, and he's making off with the scarlet heart" |
| *23* | Movie 4: ‘Hyenas’ | 00:10 - Hyena first appears in video. Narrator: "Spotted hyenas" |
| *24* | | 00:28 - Narrator: "In the outskirts of Hara in Ethiopia, two clans are coming face to face to battle over a prized resource" (Hyenas preparing for fight) |
| *25* | | 01:04 - Peak of the fight between the two hyena clans |
| *26* | | 01:27 - Narrator: "They have been fighting over access to the city" (first hyena enters city wall) |
| *27* | | 01:57 - Hyena walking up a cobble-stone street. Narrator: "And they know exactly how to get there" |
| *28* | | 02:31 - Meat market building first appears in video. Narrator: "The ancient meat market" |
| *29* | | 03:02 - Man empties bucket of meat remains on the marketplace ground. Narrator: "This tradition goes back over 400 years" |
| *30* | | 03:18 - Narrator: "They're the only animals that can. No other here has such powerful bone-crushing jaws" |

| | | |
|---|---|---|
| *31* | | 03:35 - (Sound of man hooting). Narrator: "Yousuf is calling the hyenas to his house" |
| *32* | | 04:04 - Narrator: "The inhabitants of this town believe that hyenas provide another very important service, eating the bad spirits that haunt the streets" |
| *33* | | 04:37 - Narrator: "They are perhaps the most vilified animal on our planet" |
| *34* | Story: 'Pieman' | 0:26 - "And one day I'm walking to the campus center and out comes the elusive Dean McGowan" |
| *35* | | 0:45 - "Is it true that Fordham University plans to raise tuition substantially above the inflation rate? And if so, wouldn't that be a betrayal of its mission?" |
| *36* | | 1:05 - "...which becomes the guy standing next to me mashing a cream pie into Dean McGowan's face" |
| *37* | | 1:25 - "And I say, Dean McGowan, would you care to comment on this latest attack?" |
| *38* | | 1:41 - "...and I find the editor, Jim Dwyer, who's a senior - and he will go on to win a Pulitzer Prize, that is true..." |
| *39* | | 1:57 - "So I'm banging out my story and I know it's good, and then I start to make it better" |
| *40* | | 2:09 - "Reporters call this making shit up" |
| *41* | | 2:19 - "But I had just seen the line crossed between a high-powered Dean and assault with a pastry, and I kind of liked it" |
| *42* | | 2:35 - "and I described him as a cape-wearing masked avenger. Though in fact he'd been capeless" |
| *43* | | 2:55 - "I said that he cried out, ego sum non un bestia" |
| *44* | | 3:15 - "So I finish my story, and I hand it to Dwyer, and he reads it and he says, PieMan, I love it, page one!" |
| *45* | | 3:36 - "If you want to see me again in action, be on the steps of Duane Library, Tuesday, at three o'clock. Signed, PieMan." |
| *46* | | 3:58 - "And now Sheila was different from me and all of the other Fordham students who wore flannel shirts and worked part-time jobs" |
| *47* | | 4:15 - "Although rumor had it that he got plenty of her in his office, on his desk" |
| *48* | | 4:34 - "The infamous 'No More Beer at Barbecues' rule - that's right, boo that rule" |
| *49* | | 4:57 - "...crying ego sum non un bestia. Or at least that's what it said in my story in the newspaper the next day" |
| *50* | | 5:17 - "... and that's what made him a sensation on campus. People started dressing like him and quoting him in class" |
| *51* | | 5:40 - "And now Angela and I had been flirting for two months. Or, I had been flirting with her" |
| *52* | | 5:58 - "And then I went to the bar to buy a round and I felt a tap on my shoulder" |
| *53* | | 6:16 - "And we were wondering, are you PieMan?" |
| *54* | | 6:31 - "And wasn't I really PieMan?" |
| *55* | | 6:55 - "So I looked at her and I said, Yes, Angela, I am PieMan." |

**Supplementary Table 1.** Textual reminders of the 55 sampled events. Each reminder began with a time-stamp (min:sec) referring to the onset of the event within the stimulus.
